# Identification of *Trypanosoma brucei gambiense* and *T. b. rhodesiense* in vectors using multiplexed high-resolution melt analysis

**DOI:** 10.1101/2020.04.21.052928

**Authors:** Gala Garrod, Emily R. Adams, Jessica K. Lingley, Isabel Saldanha, Stephen J. Torr, Lucas J. Cunningham

## Abstract

**Background:** Human African Trypanosomiasis (HAT) is a potentially fatal parasitic infection caused by the trypanosome sub-species *Trypanosoma brucei gambiense* and *T. b. rhodesiense* transmitted by tsetse flies. Currently, global HAT case numbers are reaching less than 1 case per 10,000 people in many disease foci. As such, there is a need for simple screening tools and strategies to replace active screening of the human population which can be maintained post-elimination for Gambian HAT and long-term Rhodesian HAT. Here we describe the development of a novel high-resolution melt assay for the xenomonitoring of *Trypanosoma brucei gambiense* and *T. b. rhodesiense* in tsetse.

**Methods:** Primers for *T. b. rhodesiense* and *T. b. gambiense* were designed to target species-specific single copy genes. An additional primer set was included in the multiplex to determine if samples have sufficient genomic material for detecting low copy number targets. The assay was evaluated on 96 wild-caught tsetse previously identified to be positive for *T. brucei s. l.* of which two were infected with *T. b. rhodesiense*.

**Results:** The assay was found to be highly specific with no cross-reactivity with non-target trypanosome species and the assay limit of detection was 10^4^ tryps/mL. HRM successfully identified three *T. b. rhodesiense* positive flies and was in agreement with the reference sub-species-specific PCRs. This assay provides an alternative to running multiple PCRs when screening for pathogenic sub-species of *T. brucei* s. l and produces results in ~2 hours, avoiding gel electrophoresis.

**Conclusions:** This method could provide a component of a simple and efficient method of screening large numbers of tsetse flies in known HAT foci or in areas at risk of recrudescence or threatened by the changing distribution of both forms of HAT.

## Introduction

Human African Trypanosomiasis (HAT) is a potentially fatal disease caused subspecies of *Trypanosoma* brucei transmitted by the bite of an infected tsetse fly (*Glossina* spp). HAT consists of two forms of the disease, each with its own distinct parasite, vectors, disease pathology, treatment and geographical distribution. Gambian HAT (gHAT),caused by *Trypanosoma brucei gambiense,* is a largely anthroponotic disease found across central and west Africa and accounts for the large majority of HAT cases (>97%) (1). GHAT can remain asymptomatic for months to years with symptoms often presenting once the infection has significantly advanced. Conversely, Rhodesian HAT (rHAT), caused by *Trypanosoma brucei rhodesiense,* is a zoonosis with occasional human infection, and represents less than 3% of all HAT cases. The World Health Organisation has targeted the elimination of HAT as a public health problem by 2020, defined as less than 1 new case per 10, 000 inhabitants in at least 90% of endemic foci and fewer than 2000 cases reported globally. Due to the zoonotic nature of rHAT, this WHO target is applicable to gHAT only. With gHAT on the brink of elimination and rHAT control remaining an important priority both in terms of human (2) and animal health, it is crucial to identify any remaining active cases, foci of transmission and areas of resurgence. The epidemiology of the two forms of HAT differ greatly therefore the monitoring and screening strategies for each form differ accordingly. Monitoring gHAT is largely reliant on the screening of the at-risk human population and treatment of cases (3). Accurate estimates of disease prevalence require high rates of coverage, which can be difficult to achieve, particularly in areas affected by conflict and political instability (4) and as the prevalence approaches <1 case per 10,000. As a result, there has been emphasis on the development of rapid diagnostic tests (RDTs) and field-friendly screening tools. In comparison to gHAT, there has been little progress or investment into the development of a screening tool for rHAT with current emphasis on passive case detection and control of the vector population. A reliance on passive detection results in a delay in the identification and treatment of infected individuals, both of which are crucial for the control of disease transmission.

With declining numbers of cases, active screening programmes are no longer cost-effective (5) and there is a need for a monitoring tool which can be maintained sustainably for rHAT and post-elimination for gHAT.

Xenomonitoring, the screening of vectors for the presence of parasites, provides a potential alternative to host sampling. This method has already been successfully utilised as a surveillance tool within the Lymphatic Filariasis elimination programme (6–8). Vectors are often routinely collected as part of vector control programmes and are far simpler and less costly to sample than either human or animal populations. Additionally, screening vectors for infection is often less time consuming and with efficient processing, can provide a view of disease transmission in real-time. Microscopy has been traditionally used for vector screening due to its low cost, high specificity and ease of use in-field. However, the sensitivity of microscopy is highly variable and morphological identification of *T. brucei gambiense* and *T. b. rhodesiense* trypanosomes is not possible.

The development of molecular tools has provided highly sensitive alternatives to traditional screening methods. PCR is widely used for the detection of *Trypanosoma* DNA, with highly sensitive assays developed for *T. brucei s. l* (9) which includes *T. b. gambiense* and *T. b. rhodesiense* along with the animal trypanosome *T. b. brucei*. Successful amplification of the target indicates the presence of one of the members of *T. brucei* s*. l.* but does not identify which sub-species is present. Identification of *T. b. gambiense* and *T. b. rhodesiense* is reliant on the detection of specific single-copy genes for each subspecies. For detection of these genes, the presence of sufficient genetic material is crucial. A negative result may indicate the absence of the target species or simply that insufficient DNA is present. To differentiate between these two scenarios, primers have been designed to screen for other single-copy genes (10), namely a single-copy phospholipase-C (GPI-PLC) expressed by all members of T*. brucei s. l.* (11–14). Samples found to have sufficient DNA can then be screened using primers specific for *T. brucei s. l.* subspecies

High-resolution melt analysis (HRM) is a post-qPCR analysis method which can be used to detect heterogeneity within nucleotide sequences. A fluorescent dye is added to the PCR reaction which intercalates into double stranded DNA. Following amplification, the amplicon is heated gradually causing the strands to separate. Separation of the double strands releases the incorporated dye causing a drop in fluorescence. The rate of DNA strand disassociation and the temperature at which it separates (Tm) is dependent on the nucleotide sequence. Different sequences will have different melting temperatures which can be used as a diagnostic identifier. HRM is a closed tube process resulting in a reduced risk of contamination and produces results in approximately two hours making circumventing gel electrophoresis, making it a faster, more specific alternative to traditional PCR. Multiplexing allows for the screening of a number of targets simultaneously, making sample processing more efficient. Here, we describe the design of a multiplexed HRM assay for the identification of the two sub-species of HAT: *T. b. rhodesiense* and *T. b. gambiense* and sufficient DNA for single-copy gene identification.

## Methods

Primers were designed to produce an amplicon of 150-350 base pairs with distinct melt temperatures. *T. b. rhodesiense* primers were derived from the sequence for the *T. brucei rhodesiense* serum-resistance-associated (SRA) protein gene (accession number AF097331.1). Primers for *T. b. gambiense* were previously designed and published by Radwanska *et al.* (15) and target the sub-species-specific glycoprotein (TgsGP: accession number AJ277951). A third primer set was designed to identify the presence of sufficient genetic material. These primers amplify a single-copy phospholipase-C (GPI-PLC) gene expressed by all members of the Trypanozoon group (11–14).

### HRM assay

HRM reactions were run in a total volume of 12.5µl consisting of 2.5µl DNA template, 6.25 µl HRM Master Mix (Thermo-start ABgene, Rochester, New York, USA), 3.25 µl sterile DNase/RNase free water (Sigma, ST. Louis, USA) and 400nM of all forward and reverse primers. Reactions were carried out on a Rotor-Gene 6000 real-time PCR machine (Qiagen RGQ system). The following protocol was followed: denaturation at 95°C for 5 minutes followed by 40 cycles and denaturation for 10 seconds at 95°C per cycle, annealing and extension for 30 seconds at 58°C, and final extension for 30 seconds at 72°C. The melting step ran from 75°C to 95°C with a temperature increase of 0.1°C every 2 seconds.

### Specificity and analytical sensitivity

The specificity of the multiplexed assay was evaluated using DNA from a range of non-target trypanosome species: *Trypanosoma congolense* Savannah (Gam2), *T. congolense* Forest (ANR3), *T. congolense* Kilifi (WG84), *T. simiae* (TV008), *T. godfreyi* (Ken7), *T. vivax* (Y486), *T. grayi* (ANR4) and *T. brucei brucei* (M249). The analytical sensitivity of the assay was assessed using a ten-fold dilution series of target species DNA. Sub-species-specific PCRs were run alongside for direct comparison of sensitivities (10,15).

### Screening of field samples

A subsample of 96 wild-caught tsetse *(Glossina swynnertoni*, *G. pallidipes*), previously shown to be *T. brucei s. l.* positive were screened individually for infection using HRM. The flies were from a collection of 5986 tsetse originally captured in 2015-2016, using odour-baited Nzi traps (16) deployed at sites in Grumeti and Ikorongo wildlife reserve, and Serengeti National Park of Tanzania. Captured flies were stored individually in 100% ethanol at room temperature and returned to LSTM for analysis. DNA extraction was carried out using Genejet DNA purification kit (Thermo K0721) according to the manufacturer’s instructions. Flies were screened using TBR PCR (9) to identify those infected with *T. brucei* s. l parasites. All *T. brucei* s. l positive flies were screened for the presence of human pathogenic trypanosomes using the multiplexed HRM. Samples were classified as positive by the presence of a peak occurring at the predicted Tm with a height above 10% of the maximum dF/dT of the highest peak. Confirmatory testing was done by processing all flies using SRA PCR for *T.b. rhodesiense* (10) and TgsGP PCR for *T. b. gambiense* (15). Two of the 96 samples were previously identified as being *T. b. rhodesiense* using SRA PCR. Further details of the collection, DNA extraction and analyses of the tsetse are reported by Lord *et al* (16).

### Results

One pair of primers per target was selected based on amplification, distinct Tm and peak fluorescence. Product sizes for each amplicon ranged from 134-319 base pairs (Table 1) with peak temperatures ranging from 79.2°C to 87.5°C.To allow for automated calling of peaks, bin widths of 1.5°C (0.75°C either side of diagnostic Tm) were set for each target.

**Table 1.**
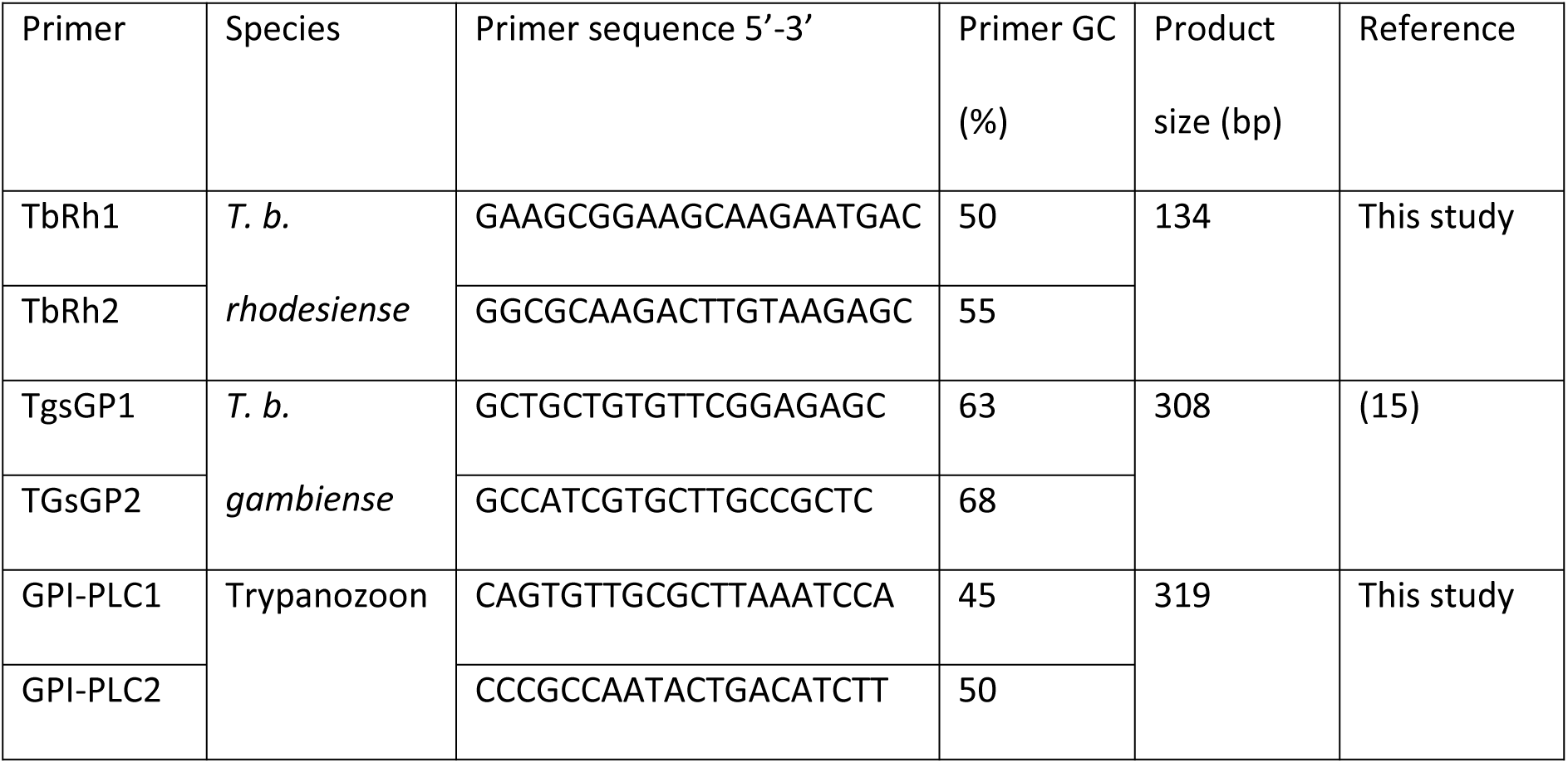
Primers included in the HRM multiplex.

### Analytical specificity and sensitivity

No non-specific amplification was seen when the assay was challenged with a range of non-target trypanosome species, namely *T. congolense* (Savannah, Kilifi and Forest subgroups), *T. vivax*, *T. simiae*, *T. simiae* Tsavo, *T. godfreyi* and *T. grayi.* The limit of detection was found to be 10^4^ trypanosomes/mL for *T. b. gambiense* and *T. b. rhodesiense* using purified DNA. When the analytical sensitivity of the HRM was compared to TgsGP and SRA PCR, the HRM was as sensitive at detecting *T. b. gambiense* and tenfold more sensitive for *T. b. rhodesiense*.

### Screening of field samples

GPI-PLC positive control peaks were produced by 43 samples (45%), indicating sufficient DNA quantity for species specific gene detection (Table 2). Of these 43 flies, three were identified as positive for *T. b. rhodesiense* DNA (**Error! Reference source not found.**). No samples were positive for *T. b. gambiense.* The HRM results were consistent with those produced by SRA and TgsGP PCR. Flies that were positive for GPI-PLC but negative for *T. b. gambiense* or *T. b. rhodesiense* were considered to be infected with livestock trypanosome *T. b. brucei*.

**Table 2.** Number of positive flies for T. b. gambiense, T. b. rhodesiense and GPI-PLC.

| Target | Positive samples |  |
| --- | --- | --- |
|  | HRM (n) | PCR (n) |
| *T. b. gambiense* | 0 | 0 |
| *T. b. rhodesiense* | 3 | 3 |
| GPI-PLC | 43 | 19 |

**Figure 1.**
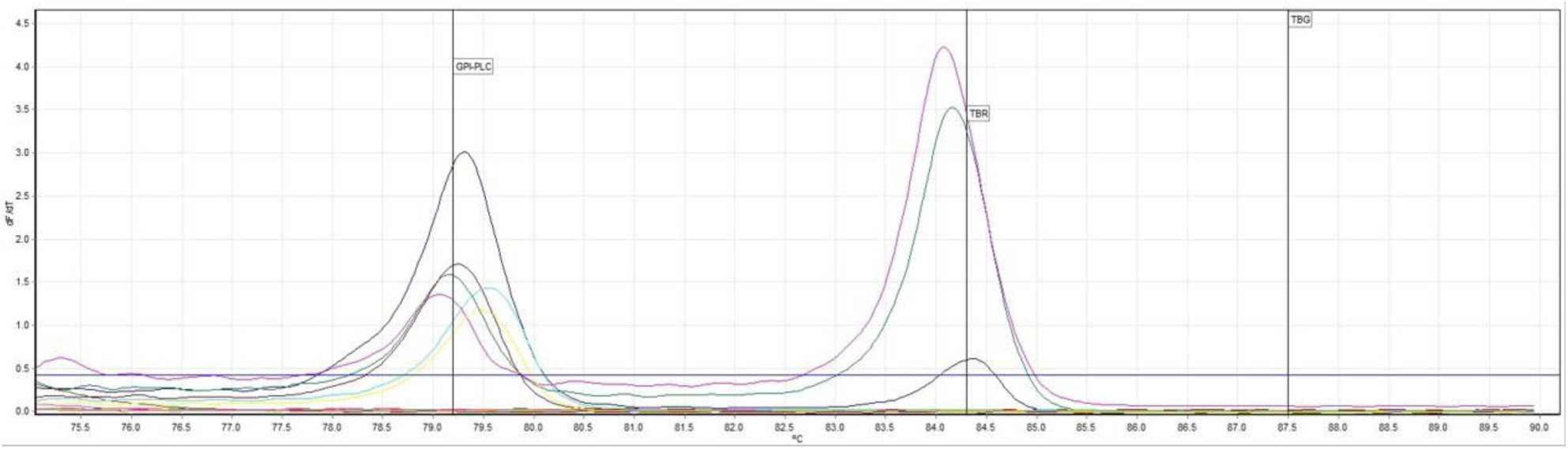
Melt profile of field samples showing three T. b. rhodesiense positives

## Discussion

Here we describe a novel high-resolution melt analysis for the detection and differentiation of *T. b. rhodesiense* and *T. b. gambiense*. This multiplexed assay screens for both pathogenic trypanosome species simultaneously with the addition of a third primer set to identify the presence of sufficient DNA to detect single-copy genes, acting as a positive control. The assay demonstrated high specificity with no cross-reaction with other non-target trypanosome species also transmitted by tsetse. The limit of detection of the HRM was lower than those reported in the literature for TgsGP (15) and SRA PCR (10). However, when the three assays were tested on a dilution series of DNA, HRM was found to be as sensitive as TgsGP PCR and 10-fold more sensitive than SRA PCR. The assay correctly identified three wild caught tsetse flies to be positive for *T. b. rhodesiense* DNA. These data were in agreement with reference sub-species PCRs. Of the 96 flies screened, 45% were found to have adequate genetic material for single-copy gene detection. As a result, 55% of tsetse remained unidentified to sub-species level. The method has three advantages over traditional PCR methods. First, the HRM time to result of ≤2 hours is faster than PCR followed by gel electrophoresis which can take over ≥3 hours for product amplification and visualisation. Second, this is a closed-tube assay which reduces contamination risk. Finally, it does not require interpretation of gel electrophoresis results. Through the use of detection bins, sample processing can also be automated, further speeding up and simplifying data analysis. The simple and fast nature of this method indicates it could be suitable for the high-throughput processing of tsetse. With prevalence of *T. b. gambiense* in tsetse from HAT foci predicted to be as low as 1 in 10^5^ (17), there is a need for a xenomonitoring tool which can be applied to large numbers of samples. With further optimisation of the assay and DNA extraction protocol, our method could be applied in the remote and low-resource settings typical of most HAT cases. This method could therefore provide the basis of a real-time trypanosome transmission monitoring platform, enabling timely reactive measures by disease control programmes. Furthermore, with the traditionally distinct geographical distributions of both Rhodesian and Gambian HAT changing due to the movement of livestock (18), human migration (19) and climate change (20–23), it may become increasingly important to simultaneously screen for both trypanosome species. The HRM allows for this and removes the risk of presumptive screening based on historic disease distributions.

The authors acknowledge that the reliance of low copy genes for target identification is a limiting factor of the described diagnostic assay. However, at present these single copy genes are the only identifiers of members of the *T. brucei s. l* and so poses a challenge to any diagnostic method based on these targets. This study was also challenged by the unavailability of any field-caught *T. b. gambiense* infected tsetse flies. As previously mentioned, trypanosome prevalence in HAT foci is predicted to be very low (17), therefore making obtaining sufficient field samples for assay validation an ongoing challenge.

In summary, we describe the development of a novel HRM assay for the detection and discrimination of human African trypanosomes in tsetse flies. The assay also incorporates an internal control, identifying samples with sufficient genomic material. The closed tube nature of the assay in addition to the relatively fast time to result lends itself to use in high-throughput xenomonitoring surveillance campaigns for HAT.

## Acknowledgments

We thank Drs Liam Morrison and Jennifer Lord for providing the Tanzanian tsetse samples and screening data collected under the auspices of the Zoonosis and Emerging and Livestock Systems (ZELS) programme (BB/L019035/1).

